# Whole-genome duplication shapes the aneuploidy landscape of human cancers

**DOI:** 10.1101/2021.05.05.442712

**Authors:** Kavya Prasad, Mathew Bloomfield, Hagai Levi, Kristina Keuper, Sara V. Bernhard, Nicolaas C. Baudoin, Gil Leor, Maybelline Giam, Cheng Kit Wong, Giulia Rancati, Zuzana Storchova, Daniela Cimini, Uri Ben-David

## Abstract

Aneuploidy – a hallmark of cancer – has tissue-specific recurrence patterns suggesting it plays a driving role in cancer initiation and progression. However, the contribution of aneuploidy to tumorigenesis depends on the cellular and genomic context in which it arises. Whole-genome duplication (WGD) is a common macro-evolutionary event that occurs in >25% of human tumors during the early stages of tumorigenesis. Although tumors that have undergone WGD are reported to be more permissive to aneuploidy than tumors that have not, it remains unknown whether WGD affects aneuploidy recurrence patterns in human cancers. Here we analyzed clinical tumor samples from 449 WGD- tumors and 157 WGD+ tumors across 22 tumor types. We found distinct recurrence patterns of aneuploidy in WGD- and WGD+ tumors. The relative prevalence of recurrent aneuploidies decreased in WGD+ tumors, in line with increased aneuploidy tolerance. Moreover, the genetic interactions between chromosome arms differed between WGD- and WGD+ tumors, giving rise to distinct co-occurrence and mutual exclusivity aneuploidy patterns. The proportion of whole-chromosome aneuploidy vs. arm-level aneuploidy was significantly higher in WGD+ tumors, indicating distinct dominant mechanisms for aneuploidy formation in WGD- and WGD+ tumors. Human cancer cell lines successfully reproduced these WGD/aneuploidy interactions, confirming the relevance of studying this phenomenon in culture. Lastly, we induced WGD in human colon cancer cell lines, and followed aneuploidy formation in the isogenic WGD+/WGD-cells under standard or selective conditions. These experiments validated key findings from the clinical tumor analysis, and revealed a causal link between WGD and altered aneuploidy landscapes. We conclude that WGD shapes the aneuploidy landscape of human tumors, and propose that the interaction between WGD and aneuploidy is a major contributor to tumor evolution.

## Introduction

Whole-genome duplication (WGD) is a macro-evolutionary genetic alteration that affects over a quarter of human tumors (Zack et al. 2013; Bielski et al. 2018). WGD promotes tumorigenesis by propagating further genomic instability, thereby creating a diverse substrate for tumor evolution (Fujiwara et al. 2005; Watkins et al. 2020); attenuating selection against mutations in essential genes (López et al. 2020); and increasing the tolerance for chromosome mis-segregation (Dewhurst et al. 2014; Kuznetsova et al. 2015). WGD has been associated with several molecular and clinical tumor features, including *TP53* mutations, higher mutational burden, increased proliferation signatures, and worse overall survival (Bielski et al. 2018; Quinton et al. 2021). While promoting tumorigenesis, WGD has also been associated with altered cellular vulnerabilities in cancer cells, such as increased sensitivity to the inhibition of the mitotic kinesin KIF18A (Quinton et al. 2021; Cohen-Sharir et al. 2021; Marquis et al. 2020).

Aneuploidy, defined as copy number alterations of whole chromosomes or chromosome arms, is a hallmark of cancer, whose contribution to cancer development and progression greatly depends on the cellular and genomic context (Ben-David and Amon 2019). Aneuploidy patterns are shaped by tumor type (Zack et al. 2013; Taylor et al. 2018; Ben-David et al. 2017), the tumors’ active oncogenic pathways (Gatza et al. 2014), and even the specific driver mutations (Ben-David et al. 2016). Moreover, aneuploidies genetically interact, such that pairs of aneuploidies can sometimes co-occur or be mutually-exclusive with one another (Ozery-Flato et al. 2011; Ravichandran et al. 2018). Tumors that have undergone WGD (hereinafter referred to as WGD+ tumors) present an elevated degree of aneuploidy, and of chromosome losses in particular, in comparison to tumors that have not undergone WGD (hereinafter referred to as WGD- tumors) (Taylor et al. 2018; Bielski et al. 2018; Shukla et al. 2020; Watkins et al. 2020; Cohen-Sharir et al. 2021). Importantly, however, whether and how WGD contributes to shaping the aneuploidy patterns of human cancers remains unknown.

In yeast, polyploidy has been associated with genomic instability and with unique cellular vulnerabilities (Storchová et al. 2006), as well as with a greater ability to explore genotypic and phenotypic solutions to stressful culture conditions in *in vitro* evolution experiments (Scott et al. 2017). Moreover, specific aneuploidies were shown to confer fitness advantage in a ploidy-dependent manner (Selmecki et al. 2015), and the adaptive value of specific chromosome-chromosome genetic interactions depended on the ploidy of the organism (Ravichandran et al. 2018). Induction of certain chromosome losses in mouse embryonic fibroblasts (MEFs) could drive tumorigenesis *in vivo* on a tetraploid, but not diploid, genomic background (Thomas et al. 2018). These studies raise the possibility that WGD may also alter the adaptive value of specific aneuploidies, and their genetic interactions with other aneuploidies, in human tumors, potentially resulting in distinct aneuploidy recurrence patterns between WGD- and WGD+ tumors.

Here, we compared the aneuploidy landscapes between WGD- and WGD+ tumors across all The Cancer Genome Atlas (TCGA) tumor types. We identified distinct aneuploidy recurrence patterns and chromosome-arm genetic interactions in WGD- vs. WGD+ tumors, then validated these results in human cancer cell lines. Furthermore, we studied the relationship between WGD and aneuploidy using genetically-matched systems of human colon cancer cell lines before and after WGD (Kuznetsova et al. 2015; Tan et al. 2019; Bloomfield et al. 2021). Our findings suggest that WGD is a major determinant of aneuploidy evolution in human cancer.

## Results

### The prevalence and general features of aneuploidy differ between WGD- and WGD+ tumors

To compare the aneuploidy landscape of WGD- vs. WGD+ tumors, we compiled genomic data from TCGA. We determined WGD status and chromosome-arm copy number alterations per sample, as previously described (Taylor et al. 2018; Cohen-Sharir et al. 2021). Aneuploidy was defined as deviation from a diploid karyotype for WGD- tumors and as deviation from a tetraploid karyotype for WGD+ tumors, and the number of whole-chromosome and chromosome-arm gains and losses were computed for each tumor sample. We focused our downstream analyses on the 22 tumor types with >20 samples in each of the WGD groups (**Supplementary Table 1**).

Across tumor types, WGD+ tumors were significantly more aneuploid than WGD- tumors (**Fig. 1a** and **Supplementary Fig. 1a**). In line with the role proposed for WGD in “buffering” the cellular consequences of chromosome-arm losses (Ganem et al. 2007), and as previously reported (Zack et al. 2013; Taylor et al. 2018; Bielski et al. 2018), the fraction of chromosome-arm and whole-chromosome losses out of all aneuploidies was significantly higher in WGD+ tumors (**Fig. 1b**). The increased aneuploidy levels of WGD+ tumors might stem from an increased prevalence of the most recurrent aneuploidies, or from an elevated tolerance to aneuploidy in general. To distinguish between these two options, we compared the relative prevalence of each aneuploidy between WGD- and WGD+ tumors. For both gains and losses, although the absolute prevalence of most recurrent aneuploidies was higher in the WGD+ tumors in all tumor types, their relative prevalence was significantly lower than in WGD- tumors (**Fig. 1c**). Moreover, in line with WGD+ tumors being more permissive to aneuploidy in general, the distribution of aneuploidy across the genome was more uniform in WGD+ than in WGD- tumors (**Fig. 1d,e** and **Supplementary Fig. 1b,c**), as measured by quantifying the deviation of the aneuploidy distributions from uniform distributions (**Methods**). Together, these data suggest that aneuploidy is not only more pervasive in WGD+ than in WGD- tumors, but also that the aneuploidy landscapes of WGD+ tumors are more promiscuous.

**Figure 1:**
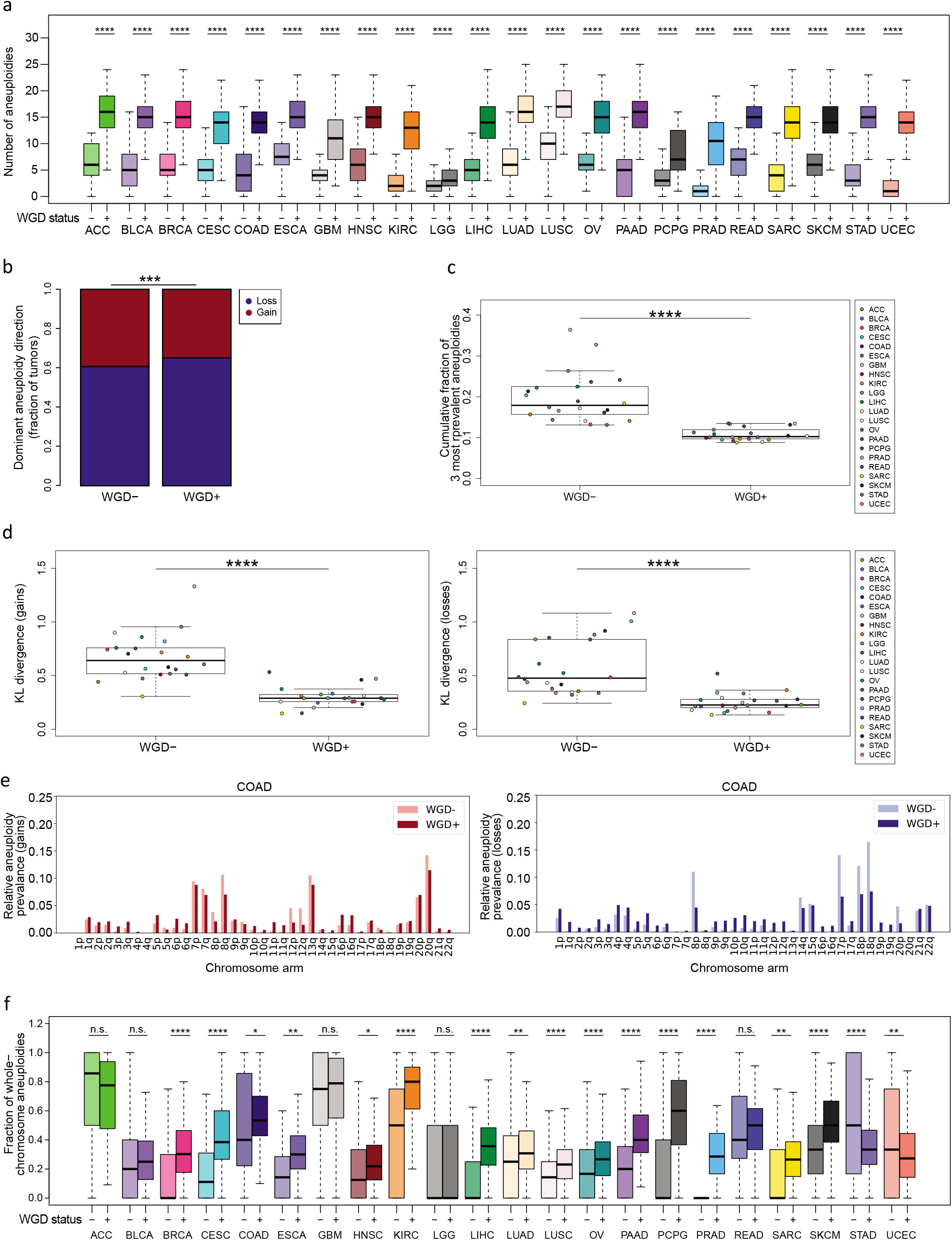
Distinct prevalence and features of aneuploidy in WGD- and WGD+ tumors. (**a**) Comparison of the number of aneuploidies between WGD- and WGD+ tumors within 22 tumor types. *, p<0.05; **, p<0.01; ***, p<0.001; ****, p<0.0001; two-tailed Student’s t-test. (**b**) Comparison of the dominant whole-chromosome aneuploidy direction (i.e., the relative prevalence of gains vs. losses) between WGD- and WGD+ tumors, across all cancer types combined. p= 2e-04; two-tailed Fisher’s Exact test. (**c**) Comparison of the cumulative fraction of the top three aneuploidies (out of all aneuploidies) between the WGD- and WGD+ tumors, across the 22 tumor types. P=6e-8; two-tailed Student’s t-test. (**d**) Comparison of the deviation of chromosome-arm gains (left) and losses (right) from a uniform distribution, between WGD- and WGD+ tumors, across the 22 tumor types. ****, p=8e-08 and p=4e-07, for gains and losses, respectively; two-tailed paired Student’s paired t-test. (**e**) The relative prevalence of chromosome-arm gains (left) and losses (right) in WGD- and WGD+ colon adenocarcinoma (COAD) tumors. (**f**) Comparison of the fraction of whole-chromosome aneuploidies (out of all whole-chromosome and arm-level aneuploidies) between WGD- and WGD+ tumors within 22 tumor types. *, p<0.05; **, p<0.01; ***, p<0.001; ****, p<0.0001; two-tailed Student’s t-test. ACC, adrenocortical carcinoma; BLCA, bladder urothelial carcinoma; BRCA, breast invasive carcinoma; CESC, Cervical squamous cell carcinoma and endocervical adenocarcinoma; COAD, colon adenocarcinoma; ESCA, esophageal carcinoma; GBM, glioblastoma multiforme; HNSC, Head and Neck squamous cell carcinoma; KIRC, Kidney renal clear cell carcinoma; LGG, Brain lower grad glioma; LIHC, Liver hepatocellular carcinoma; LUAD, Lung adenocarcinoma; LUSC, Lung squamous cell carcinoma; OV, Ovarian serous cystadenocarcinoma; PAAD, Pancreatic adenocarcinoma; PCPG, Pheochromocytoma and Paraganglioma; PRAD, Prostate adenocarcinoma; READ, Rectum adenocarcinoma; SARC, sarcoma; SKCM, Skin Cutaneous Melanoma; STAD, Stomach adenocarcinoma; UCEC., Uterine Corpus Endometrial Carcinoma

Interestingly, in 15 of the 22 tumor types, whole-chromosome aneuploidies contributed significantly more than chromosome-arm aneuploidies to the overall aneuploidy landscape in the WGD+ tumors compared to WGD- tumors (**Fig. 1f**). This finding is consistent with a high rate of chromosome mis-segregation induced by WGD (Fujiwara et al. 2005; Watkins et al. 2020; Kuznetsova et al. 2015; Dewhurst et al. 2014; Wangsa et al. 2018), and also with an increased tolerance of WGD+ tumors to whole-chromosome aneuploidies, in comparison to WGD- tumors (Dewhurst et al. 2014).

### WGD is associated with distinct aneuploidy patterns and chromosome-arm genetic interactions

Next, we compared aneuploidy recurrence patterns between WGD- and WGD+ tumors. As expected, the relative tendency of individual chromosomes and chromosome arms to be gained or lost was similar between the WGD groups (**Fig. 2a**). However, many more aneuploidies were found to be recurrent in the WGD+ group than in the WGD- group, based on a GISTIC 2.0 analysis (Mermel et al. 2011) (**Methods**; **Fig. 2a** and **Supplementary Table 2**). Overall, we identified 399 recurrent chromosome-arm aneuploidies in WGD+ tumors across the 22 tumor types, compared to 300 recurrent events in WGD- tumors (**Supplementary Table 2**). 254 (85%) of the recurrent events in WGD- tumors were also recurrent in WGD+ tumors (**Supplementary Fig. 2a**).

**Figure 2:**
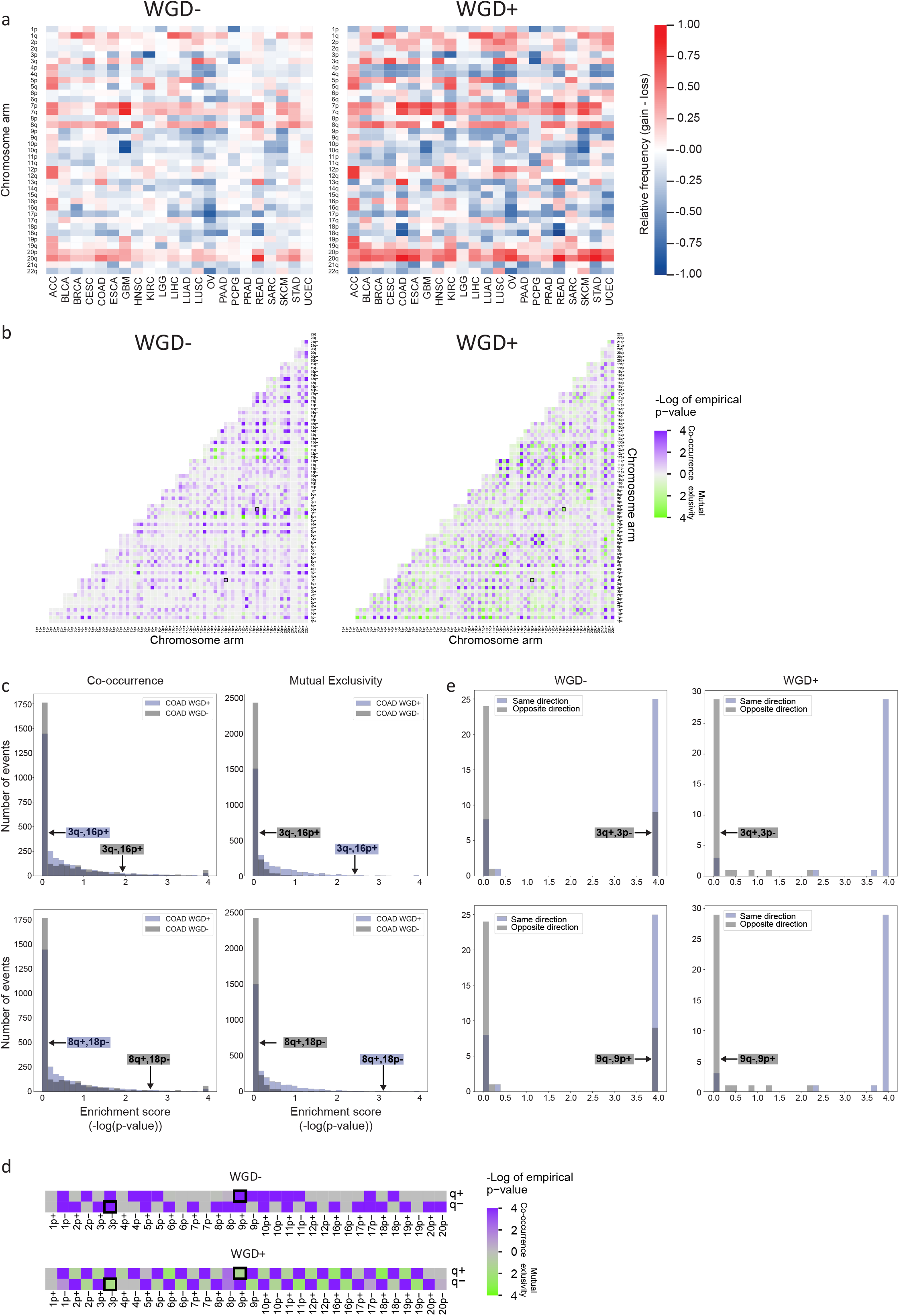
WGD is associated with significant changes in aneuploidy recurrence patterns. (**a**) Heatmaps of the relative prevalence of all chromosome-arm gains and losses in WGD- tumors (left) and WGD+ tumors (right), across 22 tumor types. The prevalence of the events was calculated by GISTIC 2.0. (**b**) Heatmaps of the significance (-log(empirical p-value)) of positive genetic interactions (co-occurrence; in purple) and negative genetic interactions (mutual-exclusivity; in green) between chromosome arms of different chromosomes in WGD- tumors (left) and WGD+ tumors (right) of colon adenocarcinomas (COAD). Note the interactions between loss of chromosome 3q (3q-) and gain of chromosome 16p (16p+), and between loss of chromosome 8q (8q-) and gain of chromosome 18p (18p+), which are significantly mutually exclusive in WGD- tumors and significantly co-occurring in WGD+ tumors (empirical p<0.05; q<0.25) and are highlighted on the heatmaps. (**c**) Histograms of the distribution of enrichment scores (defined as -log(empirical p-value)) for co-occurrence (left) and mutual exclusivity (right) of inter-chromosomal genetic interactions in WGD- vs. WGD+ COAD tumors. The discordant genetic interactions (3q-,16p+; 8q-,18p+) are highlighted on the histograms. (**d**) Heatmaps of the significance (-log(empirical p-value)) of positive genetic interactions (co-occurrence; in purple) and negative genetic interactions (mutual-exclusivity; in green) between chromosome arms of the same chromosome in WGD- tumors (left) and WGD+ tumors (right) of COAD. Note the interactions between loss of chromosome 3p (3p-) and gain of chromosome 3q (3q+), and between gain of chromosome 9p (9p+) and loss of chromosome 9q (9q-), which are significantly co-occurring in WGD- tumors and significantly mutually exclusive in WGD+ tumors (empirical p<0.05; q<0.25) and are highlighted on the heatmaps. (**e**) Histograms of the distribution of enrichment scores (defined as -log(empirical p-value)) for co-occurrence of same-direction and opposite-direction intra-chromosomal genetic interactions in WGD- (left) and WGD+ (right) COAD tumors. The discordant genetic interactions, which are co-occurring in one WGD group and mutually exclusive in the other (3p-,3q+; 9p+,9q-), are highlighted on the histograms. Tumor type abbreviations as in **Fig. 1**.

The complex karyotypes of tumors are shaped by selective pressures that can drive the gain or loss of individual chromosomes, as well as selective pressures that promote or suppress genetic interactions between chromosomes (Ozery-Flato et al. 2011; Ravichandran et al. 2018). We therefore sought to compare the chromosome-arm genetic interactions – namely, the co-occurrence and mutual-exclusivity aneuploidy patterns – between WGD- and WGD+ tumors. However, given the overall higher degree of aneuploidy in the WGD+, it is expected to observe more chromosome-arm genetic interactions in the WGD+ group, potentially confounding the comparison between the groups. Indeed, the correlation between copy number alterations (CNAs) in cancer genomes was previously shown to be confounded by the overall genomic disruption of the samples (Zack et al. 2013). To overcome this challenge, we applied a simulated annealing approach (Zack et al. 2013; Kirkpatrick et al. 1983). For each tumor type and WGD group (i.e., WGD+ and WGD-), we created 10,000 permuted matrices, maintaining both the chromosome-arm aneuploidy prevalence across tumor samples and the overall aneuploidy levels within tumor samples. We then used a hypergeometric test to determine the co-occurrence of all possible pairs of chromosome-arm aneuploidies in each tumor type, and compared the observed co-occurrence to that calculated on the permuted matrices, thus obtaining an empirical p-value for each genetic interaction (**Methods**).

Across cancer types, we found distinct chromosome-arm genetic interactions between WGD- and WGD+ tumors (**Fig. 2b**, **Supplementary Fig. 2b** and **Supplementary Table 3**). Excluding interactions between chromosome-arms of the same chromosome, a median of 78% of the significant co-occurring aneuploidy pairs in WGD- tumors were not significant in WGD+, and 80% of the significant co-occurring aneuploidy pairs in WGD+ were not significant in WGD- (**Supplementary Table 2** and **Supplementary Fig. 2c**). Surprisingly, we identified 8 events that, within a given tumor type, were significantly co-occurring in WGD- samples, but were significantly mutually-exclusive in the WGD+ samples: for example, loss of chromosome-arm 3q (Chr3q) and gain of chromosome-arm 16p (Chr16p) frequently cooccurred in WGD- colon cancer samples, but were mutually-exclusive in WGD+ colon cancers samples (**Fig. 2b,c**). Other such events are gain of Chr8q and loss of Chr18p in colon cancer, losses of Chr5q and Chr9p in breast cancer, and losses of Chr4p and Chr14q in lung adenocarcinoma (**Fig. 2c** and **Supplementary Fig. 2b,d**). We also found two events that were significantly co-occurring in WGD+ tumors, but were significantly mutually-exclusive in the WGD- tumors: loss of Chr10q and loss of Chr18p in low-grade glioma, and loss of Chr1p and gain of Chr5p in glioblastoma (**Supplementary Fig. 2b,d**). We conclude that opposite genetic interactions between specific chromosome-arm aneuploidies can exist in WGD- and WGD+ tumors. To the best of our knowledge, this is the first demonstration that the fitness of specific karyotypes is altered in human cancer following WGD, consistent with previous reports in yeast (Selmecki et al. 2015; Ravichandran et al. 2018). Notably, 9 out of these 10 genetic interactions involve a chromosome-arm alteration that is significantly associated with increased aggressiveness or worse outcome in patients (Shukla et al. 2020; Vasudevan et al. 2020), highlighting the potential clinical relevance of this finding.

We next considered the intra-chromosome interactions, i.e. the interactions between the p arm and the q arm within chromosomes. We found that opposite-direction chromosome-arm aneuploidies within the same chromosome (that is, loss of the p arm and gain of the q arm, or vice versa) are much more common in WGD- tumors. Whereas such opposite-direction alterations rarely occurred (and were therefore mutually-exclusive) in WGD+ tumors, they often co-occurred in WGD- tumors (**Fig. 2d,e, Supplementary Fig. 2e** and **Supplementary Table 4**). For example, Chr3p loss and Chr3q gain are mutually-exclusive events in WGD+, but are co-occurring events in WGD- colon cancer tumors; and so are Chr9p gain and Chr9q loss (**Fig. 2d,e**). This is in line with our observation that whole-chromosome aneuploidy (relative to chromosome-arm aneuploidy) accounts for a larger fraction of the aneuploidy landscape in WGD+ than in WGD- tumors (**Fig. 1f**): when a chromosome-arm is altered in a WGD+ tumor, the reciprocal arm is likely to be altered in the same direction; in contrast, when a chromosome arm is altered in a WGD- tumor, the reciprocal arm may be altered either in the same direction or in the opposite direction (**Fig. 2d**, **Supplementary Fig. 2e** and **Supplementary Table 3**). Together, these findings suggest that the contribution of distinct mechanisms of aneuploidy formation to tumor evolution is WGD-dependent.

### Human cancer cell lines successfully recapitulate the association of WGD with aneuploidy features

Cancer cell lines are extensively used for the research of WGD and aneuploidy (Ben-David et al. 2019). We therefore wanted to examine whether the effects of WGD on the aneuploidy landscapes are evident in cell lines as well. To compare the aneuploidy landscapes between WGD- and WGD+ cancer cell lines, we compiled data from the Cancer Cell Line Encyclopedia (CCLE) (Ghandi et al. 2019) and determined their WGD status and chromosome-arm alterations, as we previously described (Cohen-Sharir et al. 2021). We focused our analyses on the 14 cancer types with >5 samples in each of the WGD groups (**Supplementary Table 5**). In agreement with our findings from the TCGA clinical samples, WGD+ cell lines had significantly more aneuploidy events than WGD- cell lines across all lineages (**Fig. 3a**), and the relative fraction of whole-chromosome aneuploidies was higher in the WGD+ tumors for 13 out of the 14 cancer types (albeit reaching statistical significance only in 4 of them; **Fig. 3b**). The same cancer-type specific recurrent aneuploidies observed in TCGA (**Fig. 2a**) were observed in the cell lines (**Fig. 3c** and **Supplementary Table 6**). Moreover, although the absolute prevalence of the most recurrent aneuploidies was higher in WGD+ than in WGD- cell lines (**Fig. 3c**), their relative prevalence was lower (**Fig. 3d**), consistent with the findings from the primary tumors. We conclude that the associations between WGD and aneuploidy features are conserved between human tumors and human cancer cell lines.

**Figure 3:**
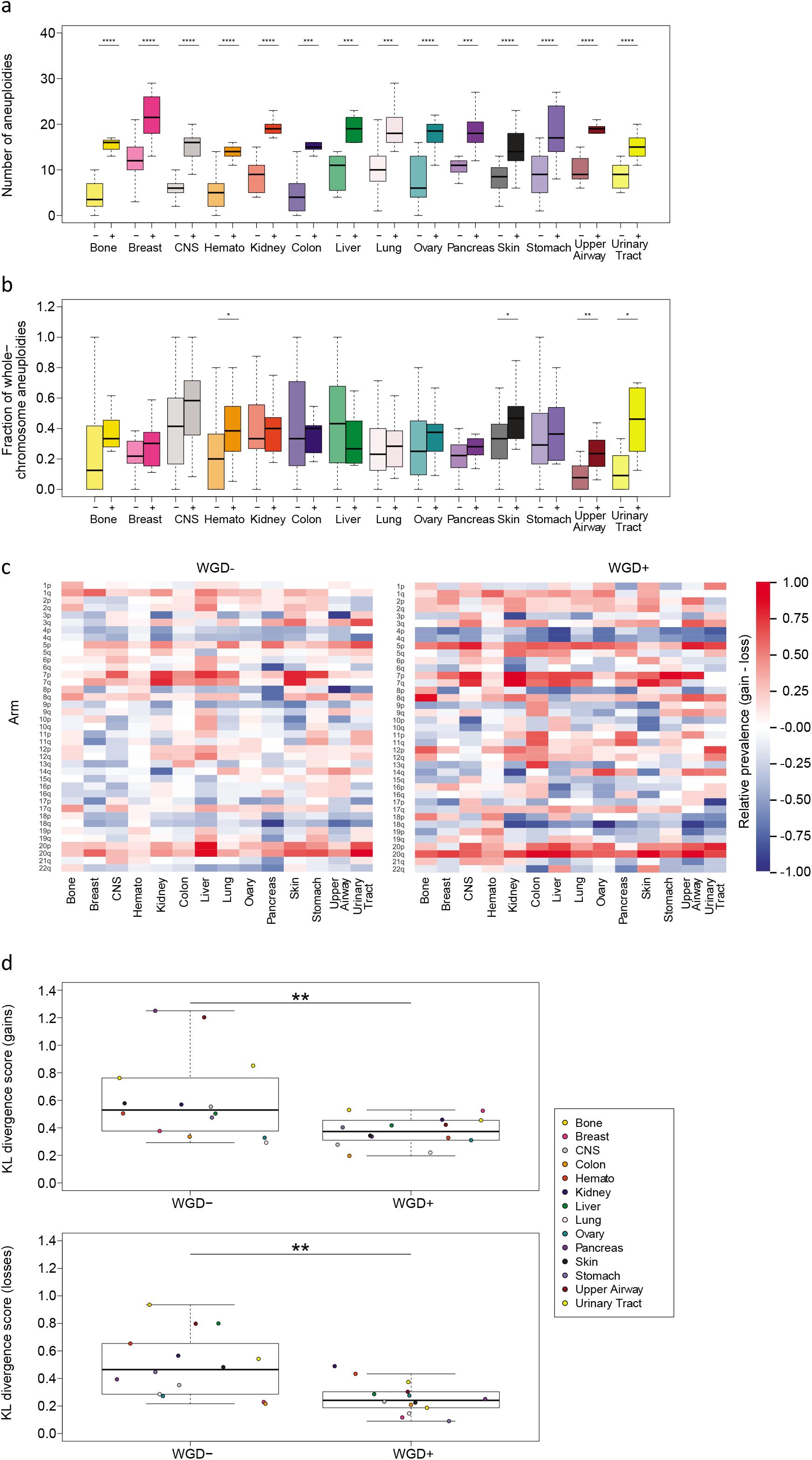
The association between WGD and aneuploidy is conserved in human cancer cell lines. (**a**) Comparison of the number of aneuploidies between WGD- and WGD+ cell lines within 14 cancer types. *, p<0.05; **, p<0.01; ***, p<0.001; ****, p<0.0001; two-tailed Student’s t-test. (**b**) Comparison of the fraction of whole-chromosome aneuploidies (out of all whole-chromosome and arm-level aneuploidies) between WGD- and WGD+ cell lines within 14 cancer types. *, p<0.05; **, p<0.01; ***, p<0.001; ****, p<0.0001; two-tailed Student’s t-test. (**c**) Heatmaps of the relative prevalence of all chromosome-arm gains and losses in WGD- cell lines (left) and WGD+ cell lines (right), across 14 cancer types. The prevalence of the events was calculated by GISTIC 2.0. (**d**) Comparison of the KL-divergence values of chromosome-arm gains (left) and losses (right) between WGD- and WGD+ cell lines, across the 14 cancer types. **, p=0.008 and p=0.001, for gains and losses, respectively; two-tailed paired Student’s paired t-test. CNS, central nervous system; Hemato, hematologic malignancies.

### Isogenic human cancer cell lines reveal a causal relationship between WGD and aneuploidy landscapes

To study the consequence of WGD in human cells, we previously developed genetically-matched systems of human colon cancer cell lines before and after WGD (Kuznetsova et al. 2015; Tan et al. 2019; Bloomfield et al. 2021). These cell lines are aneuploid, chromosomally unstable, and exhibit increased tumorigenic behavior and drug resistance (Kuznetsova et al. 2015; Tan et al. 2019; Bloomfield et al. 2021). We therefore turned to these systems to further characterize the relationship between WGD and aneuploidy in human cancer.

We first compared the near-diploid WGD- HCT116 cells to their WGD+ derivatives HPTs (HCT116 Post Tetraploid cells) (Kuznetsova et al. 2015). We used multicolor fluorescence *in situ* hybridization (mFISH) to karyotype single cells from the parental HCT116 population and from two independent HPT single cell-derived populations. In comparison to HCT116 cells, HPT cells were more aneuploid in general (**Fig. 4a**), and displayed a higher relative fraction of chromosome losses in particular (**Fig. 4b**), and whole-chromosome aneuploidies (compared to both arm-level and structural aneuploidies; **Fig. 4c**). The HPT cells remained chromosomally unstable, and did not converge on an optimal karyotype (**Fig. 4d**), in line with WGD+ cells being more tolerant to a variety of aneuploidies. We confirmed the mFISH findings using data from single-cell DNA sequencing (scDNAseq) of the HCT116 and HPT cells (Cohen-Sharir et al. 2021) (**Supplementary Fig. 3**). Importantly, we included in the mFISH analysis a population of HCT116 cells with a stable aneuploidy, which have not undergone WGD (Passerini et al. 2016). When the clonal aneuploid chromosome itself was excluded, these WGD- aneuploid HCT116 cells showed a similar degree of aneuploidy, losses/gains ratio, whole-chromosome/chromosome-arm ratio, and overall chromosomal stability, as the parental near-diploid HCT116 cells (**Fig. 4a-d**). These results confirm that it is WGD *per se* that underlies the observed changes in the aneuploidy landscape.

**Figure 4:**
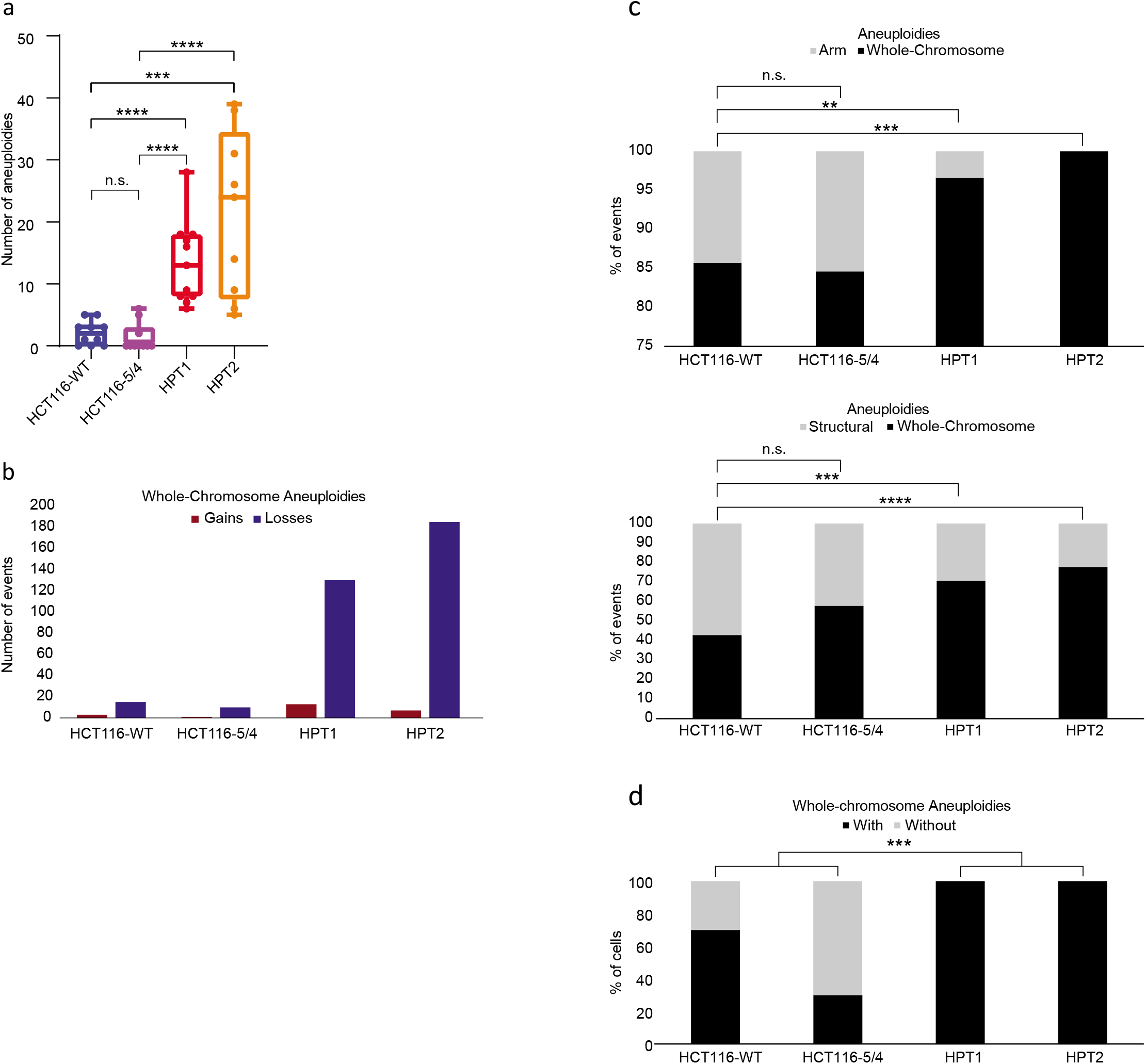
Isogenic WGD- and WGD+ HCT116 cell lines demonstrate a causal effect of WGD on aneuploidy landscapes. (**a**) mFISH-based comparison of the number of aneuploidies between the near-diploid parental HCT116 cells (HCT116-WT), HCT116 cells with two extra copies of chromosome 5 introduced through microcell-mediated chromosome transfer (HCT116-5/4) (Stingele et al. 2012), and two WGD+ HCT116 clones (HPT1 and HPT2) (Kuznetsova et al. 2015). n.s., p>0.05, ***, p<0.001; ****, p<0.0001; two-tailed Student’s t-test. (**b**) The number of whole-chromosome gains and losses observed by mFISH, in HCT116 and its derived clones (>10 single cells per clone). (**c**) The relative fraction of whole-chromosome aneuploidies relative to arm-level aneuploidies (top) or structural aneuploidies (bottom; including armlevel aneuploidies, translocations and smaller structural alterations), in HCT116 and its derived clones. n.s., p>0.05, **, p<0.01, ***, p<0.001; ****, p<0.0001; one-tailed Fisher’s Exact test. (**d**) The fraction of cells with non-clonal whole-chromosome aneuploidies, a measure of karyotypic heterogeneity, in HCT116 and its derived clones.***, p<0.001; one-tailed Fisher’s Exact test.

Lastly, we inhibited cytokinesis to induce WGD in another near-diploid WGD- human colon cancer cell line, DLD1, and isolated 11 WGD+ DLD1 clones (**Methods**). In comparison to their parental cells, WGD+ cells were generally more aneuploid (**Fig. 5a**), their aneuploidy landscape was dominated by chromosome losses (**Fig. 5b**) and whole-chromosome aberrations (**Fig. 5c**), and were more karyotypically heterogeneous (**Fig. 5d**), in complete agreement with our findings with the HCT116/HPT cells. Moreover, by inhibiting cytokinesis and comparing the karyotypic evolution of WGD- and WGD+ cells within the same culture (**Fig. 5e**), we found that aneuploidy and karyotypic heterogeneity accumulated quickly in the WGD+ cells, but not in the WGD- cells (**Fig. 5f**). 12 days of *in vitro* culture (as in Baudoin et al. 2020) were sufficient to reproduce the karyotypic features that we found to be associated with WGD+ tumors: increased levels of karyotypic heterogeneity, chromosome losses, and whole-chromosome aneuploidies, relative to WGD- cells (**Fig. 5g,h**). Interestingly, despite the selection pressure associated with single-cell cloning, no specific aneuploidy was associated with the derivation of the WGD+ clones (**Supplementary Fig. 4a**). To further study karyotypic evolution under selection, we transferred the day-12 WGD+ cells to soft-agar and karyotyped four of the macroscopic colonies emerging from them (**Supplementary Fig. 4b**). Karyotypic heterogeneity was maintained, and no recurrent aneuploidies were detected across these colonies (**Supplementary Fig. 4c**). Therefore, there was no evidence for rapid selection of a specific karyotype in either of these challenges, confirming that WGD+ cancer cells are permissive to a wide variety of aneuploidies and can find multiple evolutionary solutions to induced stress.

**Figure 5:**
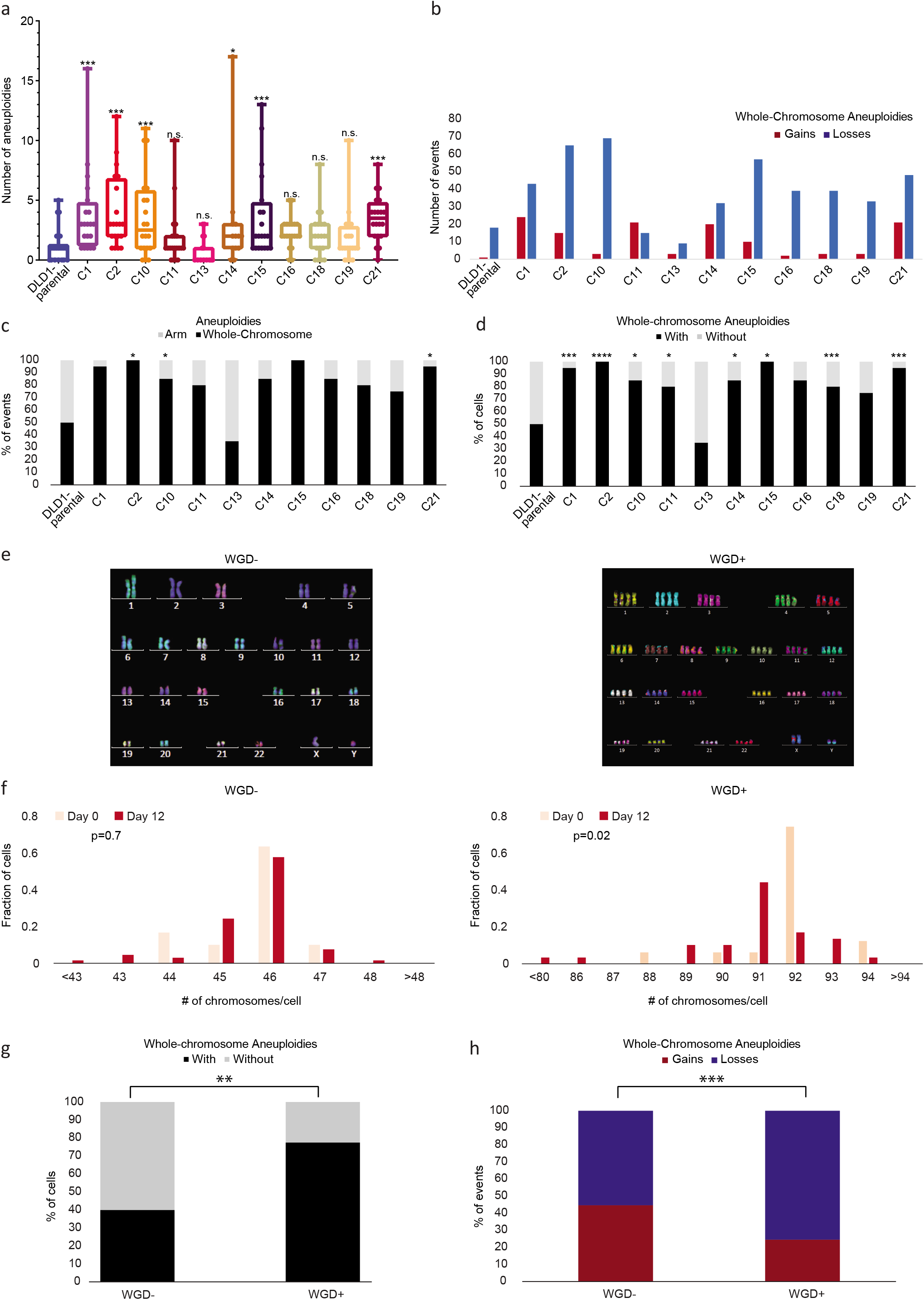
Isogenic WGD- and WGD+ DLD1 cell lines demonstrate a causal effect of WGD on aneuploidy landscapes. (**a**) mFISH-based comparison of the number of aneuploidies between near-diploid parental DLD1 cells (DLD1-parental), and 11 DLD1-derived WGD+ clones. n.s., p>0.05, *, p<0.05, ***, p<0.001; two-tailed Student’s t-test. All comparisons are with the DLD1-parental line. (**b**) The number of whole-chromosome gains and losses observed by mFISH, in DLD1 and its derived clones (>20 cells per clone). (**c**) The relative fraction of whole-chromosome aneuploidies relative to arm-level aneuploidies in DLD1 and its derived WGD+ clones. *, p<0.05; one-tailed Fisher’s Exact test. All comparisons are with the DLD1-parental line. (**d**) The fraction of cells with whole-chromosome aneuploidies, a measure of karyotypic heterogeneity, in DLD1 and its derived clones. *, p<0.05, ***, p<0.001, ****, p<0.0001; one-tailed Fisher’s Exact test. All comparisons are with the DLD1-parental line. (**e**) Representative karyograms of WGD- and WGD+ DLD1 cells taken from the same cell population right after cytokinesis inhibition. (**f**) Histograms of the chromosome count of WGD- and WGD+ cells at d0 (right after cytokinesis inhibition) and at d12 (twelve days after cytokinesis inhibition). p=0.7 and p=0.02 for the difference in the means of the distributions between d0 and d12, in the WGD- and WGD+ populations, respectively; two-tailed Mann-Whitney test. For WGD- do: means=45.67, S.D.=0.88; for WGD- d12: means=45.61, S.D.=1.01; for WGD+ d0: means=91.82, S.D.=1.33; for WGD+ d12: means=90.81; S.D.=2.10. (**g**) Comparison of the fraction of cells with whole-chromosome aneuploidies, a measure of karyotypic heterogeneity, in WGD- and WGD+ DLD1-derived cells at d12 following cytokinesis inhibition. **,p<0.01; one-tailed Fisher’s Exact test. (**h**) The fraction of whole-chromosome gains and losses in WGD- and WGD+ DLD1-derived cells at d12 following cytokinesis inhibition. ***, p<0.001; one-tailed Fisher’s Exact test.

## Discussion

In recent years, it has become clear that the evolution of tumor aneuploidy is contextdependent, and is affected by the cell lineage, tumor developmental stage, the tumor environment, and the immune system (Ben-David and Amon 2019). The genomic background from which aneuploidy arises plays an important role in determining the adaptive value of emerging chromosomal alterations (Ben-David et al. 2016; Gatza et al. 2014; Thomas et al. 2018). Here, we systematically compared the aneuploidy landscapes of WGD- and WGD+ tumors across all major tumor types, and found them to be distinct in important ways. The elevated degree of aneuploidy and increased tolerance to chromosome loss in WGD+ tumors were already reported prior to our study (Zack et al. 2013; Taylor et al. 2018; Quinton et al. 2021; Bielski et al. 2018). However, our analysis yielded several novel insights: First, we show that WGD+ tumors are indeed more tolerant to a wider range of aneuploidies, as evident by their more diverse aneuploidy patterns. Second, whole-chromosome aneuploidies constituted a larger fraction of the karyotypic alterations in WGD+ tumors, suggesting that the increased chromosome mis-segregation and/or aneuploidy tolerance resulting from WGD (Fujiwara et al. 2005; Watkins et al. 2020; Kuznetsova et al. 2015; Dewhurst et al. 2014; Wangsa et al. 2018) shape the aneuploidy landscapes of the tumors. Third, whereas tumor type-specific recurrent aneuploidies are shared by WGD- and WGD+ tumors, the genetic interactions between chromosome arms are WGD-dependent; in extreme cases, events that co-occur in WGD- tumors are mutually exclusive in WGD+ tumors, and *vice versa*. Fourth, opposite-direction chromosome-arm alterations are more common in WGD- tumors, again suggesting WGD- dependent mechanisms of aneuploidy formation and selection. Fifth, we show that the abovementioned associations are conserved in human cancer cell lines. Lastly, we use isogenic human cancer cell lines to demonstrate that WGD induction recapitulates the main changes observed in clinical samples, thereby demonstrating that the observed associations are causative rather than merely correlative. These isogenic cell line models are therefore appropriate for studying the interplay between WGD and aneuploidy in human cancer.

Ploidy-specific fitness advantage conferred by an aneuploid chromosome was previously reported in yeast (Selmecki et al. 2015). Recently, the aneuploidy landscape induced by CIN in yeast was shown to be more diverse in diploid yeast than in haploid yeast (Ravichandran et al. 2018), consistent with our findings from human tumors. Furthermore, significantly different aneuploidy recurrence patterns were observed between haploid and diploid yeast, indicating ploidy-dependent genetic interactions between specific chromosomes (Ravichandran et al. 2018). We now show that the fitness effects of specific chromosome-arm aneuploidy interactions can be ploidy-dependent in human cancer as well. Interestingly, a recent thermodynamic analysis of copy number states in human cancer revealed that WGD+ tumors are characterized by a distinct energy landscape, which is predicted to select for specific amplifications and deletions that favor the fitness of WGD+ tumors (Remacle et al. 2020). This theoretical thermodynamic analysis matches our empirical findings from tumors and cancer cell lines.

The finding that WGD+ tumors are skewed towards whole-chromosome aneuploidies suggests that chromosome mis-segregation plays a greater role in shaping the aneuploidy landscape of WGD+ tumors compared to WGD- tumors. Whether this is mostly due to increased prevalence of mis-segregation, increased tolerance for whole-chromosome aneuploidies, or both, remains an open question. It is also tempting to speculate that by comparing within-chromosome aneuploidy patterns between WGD- and WGD+ tumors, the “driver” chromosome-arm could be identified. For example, in WGD+ colon cancer, loss of chromosome 8p arises together with gain of chromosome 8q in ~20% of the tumors (**Fig. 2d**). At the same time, a gain of the entire chromosome 8 is much more prevalent than a loss of that chromosome in this tumor type (**Fig. 2d**), suggesting that it is the gain of Chr8q that is likely the driving force underlying the recurrence of this chromosome gain. This reasoning remains to be functionally tested in future studies.

Finally, the genetically-matched cell line models before and after WGD induction enabled us to move from correlation to causation, demonstrating that WGD induction indeed alters some features of tumor aneuploidy. Notably, we did not observe selection for a specific aneuploidy profile in any of our *in vitro* approaches, which included single-cell cloning and growth in soft-agar. While these findings are in line with WGD being more permissive to aneuploidy in general, this does not mean that specific karyotypes would not get selected for under different selection pressures (Rutledge et al. 2016; Ippolito et al. 2020; Lukow et al. 2020). Further studies are necessary in order to systematically map the interactions between WGD and specific aneuploidies across a variety of selection pressures.

In summary, our study shows that WGD shapes the aneuploidy landscapes of human tumors both quantitatively and qualitatively. Therefore, when examining the tumorigenic role of a specific aneuploidy or of a specific interaction between a pair of cancer aneuploidies, the WGD status of the tumor ought to be considered. More broadly, WGD should be accounted for when studying and reporting aneuploidy recurrence patterns of human cancer.

## Methods

### Data collection and preparation

The WGD status and chromosome arm aneuploidy landscapes were determined for 33 TCGA tumor types, as reported by Taylor et al. (Taylor et al. 2018). Samples of each of the 33 tumor types were separated into WGD- and WGD+ groups based on the absence or presence of genome doubling events. Only the 22 tumor types with a minimum of 20 samples in each group were retained for further analyses.

The WGD status and chromosome arm aneuploidy landscapes were determined for the CCLE cell lines, as reported in Cohen-Sharir et al. (Cohen-Sharir et al. 2021). Cell lines were classified as WGD- if their ploidy was <2.5, and were classified as WGD+ if their ploidy was >3. Only the 14 lineages with a minimum of 5 samples in each group were retained for further analyses. One stomach cancer cell line, KE97, was omitted from the analyses due to its later removal from the CCLE data set due to apparent misclassification(https://web.expasy.org/cellosaurus/CVCL_3386).

For each group, WGD- and WGD+, the number of chromosome-arm gains and chromosome-arm losses was computed per sample, and the degree of aneuploidy was determined as the number of whole-chromosome and arm-level events per sample (relative to the basal ploidy). For the calculation of the fraction of whole-chromosome aneuploidies vs. chromosome-arm aneuploidies, the acrocentric human chromosomes (13, 14, 15, 21, 22) were excluded. All analyses were performed for the TCGA and CCLE data sets separately.

### Aneuploidy recurrence analysis

Recurrent chromosome-arm aneuploidies in the WGD- and WGD- groups of each tumor type were determined by GISTIC 2.0 (Mermel et al. 2011). GISTIC 2.0.23 was applied with the hg19 build of the human genome. Segmentation files were obtained separately for each TCGA tumor type from the GDAC Firehose portal (https://gdac.broadinstitute.org/). A single segmentation file for CCLE, containing all cellline types, was downloaded from the Broad Institute CCLE portal (https://data.broadinstitute.org/ccle_legacy_data/dna_copy_number/). Each data set was split based on the WGD status of the samples, as described above. GISTIC was run with the ‘broad’ setting (using the default settings for all other parameters) in order to obtain results for each chromosome arm (as opposed to focal regions). The sex chromosomes and the short arms of acrocentric chromosomes were removed. For each WGD group of each tumor type, recurrent aneuploidies were determined as those with a prevalence threshold of >=10% of the samples and a significance threshold of q<0.05.

The positive and negative genetic interactions of all possible chromosome-arm pairs were determined, considering gains and losses of each chromosome-arm separately. First, within the WGD- and WGD+ groups of each tumor type, the chromosome-arm pairs were split into two categories – inter-chromosomal pairs and intra-chromosomal pairs – based on whether the two arms involved belonged to different chromosomes or to the same one. Chromosome-arm events whose co-prevalence was significantly higher than expected by chance (empirical p-value<0.05; empirical q-value<0.25) were defined as co-occurring. Chromosome-arm pairs whose co-prevalence was significantly lower than expected by chance (empirical p-value<0.05; empirical q-value<0.25) were defined as mutually-exclusive. A genetic interaction score (also referred to as an enrichment score) was defined as the log10(p-value) of the interaction, with cooccurring events receiving positive scores and mutually-exclusive events receiving negative scores.

### Permutation analysis

For each tumor type, the tumor samples were split into the WGD- and WGD+ groups, as described above. For each group, a matrix was created for recording the aneuploidy profiles, where rows were sample identifiers, columns were chromosome arms, and the values in each cell indicated gain (1), loss (−1) or neutral (0). To calibrate the significance of genetic interactions between WGD- and WGD+ data sets, therefore controlling for a potentially inflated rate of significant interactions in the WGD+ group, a permutation analysis was performed based on a simulated annealing approach described in Zack et al. (Zack et al. 2013). Specifically, permuted matrices were generated while maintaining the prevalence of each aneuploidy event. We repeatedly drew two samples and an aneuploidy event and swapped the values of the aneuploidy event between the samples. First, 100K such swaps were performed. Then, for additional 1M iterations, an E score was calculated as following:

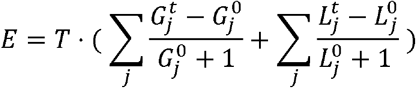

Where 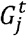 is the # of gains in sample *j*, 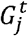 is the # of gains after iteration *t*, 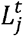 is the # of losses after iteration *t* in sample *j*, and T is the “temperature” factor that increases in each iteration 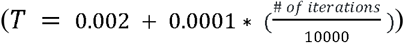; and the swap was conducted with a probability of 1 – *E*. This process was repeated 10,000 times, resulting in 10,000 permuted matrices for each WGD group in each tumor type. To enable parallelization and reproducible results, a sequence of seeds was used for the swaps: for the first 100K iterations, the sequences of seeds were unique per permuted matrix, whereas for the following 1M swaps a constant sequence of seeds was used.

Next, for each permuted matrix, the significance of co-occurring and mutually-exclusive events was calculated using a hypergeometric test, thereby generating a background distribution of 10,000 (permuted) p-values per genetic interaction. Finally, for each genetic interaction, the observed p-value was compared to the background p-value distribution, and the empirical p-value for that interaction was calculated based on its relative ranking.

### Cell lines and culture conditions

DLD1 cells (ATCC CCL-221) were obtained from the American Type Culture Collection (ATCC, Manassas, VA) and were maintained in RPMI 1640 medium with ATCC modification (Thermo Fisher Scientific-Gibco, CA, USA), supplemented with 10% fetal bovine serum (FBS; Thermo Fisher Scientific) and 1% antibiotic-antimycotic (Thermo Fisher Scientific), in a humidified incubator at 37°C and 5% CO2. Tetraploid DLD1 clones were generated by treating diploid cells with 1.5 μg/mL dihydrocytochalasin B (DCB; Sigma Aldrich, St. Louis, MO) for 20 hrs to induce cytokinesis failure. After treatment, the cells were washed four times with cell culture medium and allowed to grow an additional 1–2 days before the isolation of single cells by limiting dilution in 96-well-plates. Only wells containing a single cell were expanded into clonal cell lines and used for further experimentation after confirming ploidy by chromosome counting. For 12-day evolution experiments, DLD1 cells were treated with 1.5 μg/mL DCB for 20 hrs, followed by four washes in cell culture media. Following washout (day 0), cells were either used for chromosome spread harvesting or kept in culture with normal supplemented cell culture medium and passaged on days 2, 6 and 10 into new tissue culture flasks. On day 10, a tissue culture flask was prepared for chromosomes spread harvesting on day 12. Because <100% of the cells went through mitosis during the 20 hr treatment (Baudoin et al. 2020), both WGD- and WGD+ cells could be analyzed both at day 0 and at day 12.

HCT116 H2B-GFP cells were cultured in DMEM (Thermo Fisher Scientific), supplemented with 10% fetal bovine serum (FBS; Thermo Fisher Scientific-GIBCO), 50 IU/ml penicillin, and 50μg/mL streptomycin (Thermo Fisher Scientific), in a humidified incubator at 37°C and 5% CO2. Tetraploid HCT116 clones were previously generated, as described in Kuznetsova et al.(Kuznetsova et al. 2015), by treating diploid cells with 0.75 μg/mL dihydrocytochalasin D (DCD; Sigma Aldrich) for 18 hrs to induce cytokinesis failure. After treatment, the cells were washed, placed into a drug-free medium and subcloned by limiting dilution in 96-well plates. Only wells containing a single cell were expanded into clonal cell lines and used for further experimentation after confirming ploidy by chromosome counting.

### Chromosome spreads and chromosome counting

DLD1 cells were seeded in T-25 flasks, allowed to adhere for at least 12 hours, then treated with 50 ng/mL colcemid (Invitrogen – Karyomax, Waltham, MA) for 5 hrs. HCT116 cells were treated with 50 ng/mL colchicine (SERVA, Heidelberg, Germany) for 4.5 hrs. Cells were trypsinized, and centrifuged at 1,000 rpm for 5 min. Pre-warmed (37°C) hypotonic solution (0.075 M KCl) was added dropwise to the disrupted cell pellet and the cell suspension was incubated for 15-18 min at 37°C. Freshly prepared fixative (3:1 methanol:glacial acetic acid) was added. Chromosome spreads were dropped on wet glass slides, and either stained with 300 nM DAPI (Thermo Fisher Scientific – Invitrogen), or processed for mFISH (see next section). DAPI stained slides were mounted with antifade solution (90% glycerol, 0.5% N-propyl gallate) and sealed under 22×50 mm coverslips (Corning Incorporated, Corning, NY) with nail polish. Images of DAPI-stained chromosome spreads were acquired with a Nikon Eclipse Ti inverted microscope equipped with a ProScan automated stage (Prior Scientific, Cambridge, UK), CoolSNAP HQ2 CCD camera (Photometrics, AZ, USA), and Lumen200PRO light source (Prior Scientific, Cambridge, UK) using either a 60X/1.4 NA or 100X/1.4NA Plan-Apochromatic phase contrast objective. After image acquisition, chromosomes in individual spreads were counted using NIS elements (Nikon instruments Inc., NY, USA) or FIJI(Schindelin et al. 2012).

### Multicolor fluorescence in situ hybridization (mFISH)

For HCT116 cells, mFISH was performed with a DNA probe mixture (24XCyte Human Multicolor FISH Probe Kit, MetaSystems) as previously described (Kuznetsova et al. 2015). Briefly, fixed cells were dropped on a slide, pepsinized, incubated in a coplin jar for 2 min at 37°C and rinsed twice for 7 min each time in 1xPBS. Subsequently, the slides were dehydrated in EtOH series, 3 min each, and baked at 61.4°C for 1 h. Denaturation at 72°C for 1min 30 sec in 70% Formamide/2xSSC, and dehydration in EtOH series was followed by *in situ* hybridization. Here, 4 μl of denatured, preannealed probe was applied on a slide, covered with a coverslip, sealed with rubber cement and incubated in a dark chamber overnight at 37°C. Subsequently, rubber cement was removed, coverslip was removed by soaking the slide in 4XSSCT and the slides were washed 3×5min in 0.1xSSC at 62°C. For biotin detection, additional steps with streptavidin-Alexa Fluor 488 (Molecular Probes) were performed. Finally, the slides were washed twice each time 7 min in 4xSSCT at 42°C, mounted in Vectashield Antifade solution (4’,6 diamidino-2-phenylindole, Vector laboratories H-1200, Axxora/Alexis, Lörrach, Germany) with DAPI and cover with 24×60 mm coverslip sealed with a nail polish. The spreads were manually analyzed on the Zeiss Observer.Z1 microscope, Plan Apochromat 63x magnification oil objective in DAPI, CFP, GFP, Cy3, Texas Red and Cy5 channels.

For DLD1 cells, mFISH was performed with 24XCyte human probe cocktail (MetaSystems) according to manufacturer’s protocol. Briefly, chromosome spreads were incubated in 2X saline-sodium citrate (SSC) buffer at 70°C for 30 min, cooled to room temp for 20 mins, and washed for 1min each in 0.1X SSC and 0.07M NaOH at room temperature. Next, slides were washed at 4°C in 0.1X SSC for 1 min, followed by 2X SSC for 1 min, then 70% ethanol, 90% ethanol, and 100% ethanol for 1 min each. Probes were added to the slide, sealed under a coverslip with rubber cement, and hybridized overnight at 37°C. The coverslip was then removed, and slides were washed for 2 min in 0.4X SSC at 70°C and for 30 sec in 2XSCC, followed by a quick rinse in DIH_2_O and mounting in DAPI/Antifade counterstain buffer (MetaSystems) for imaging. Images were acquired with a Zeiss Axioplan 2 motorized microscope (Carl Zeiss, Inc., Oberkochen, Germany) equipped with an Ikaros4/Isis4 system (MetaSystems). Analysis was performed using Isis 4 software.

### Single cell DNA sequencing (scDNAseq)

Single cell DNA sequencing data of HCT116 and HPT cells were obtained and processed as described in Cohen-Sharir et al. (Cohen-Sharir et al. 2021) For each single cell, the number of whole-chromosome, chromosome-arm and sub-chromosomal (structural) copy number changes were extracted from the AneuFinder plots (Bakker et al. 2016).

### Soft agar assay

Cells were grown in T-25 flasks. On the day of experimental set up, melted 2% agar (Fisher Scientific, NJ, USA) was mixed 1:1 with 2X DMEM supplemented with 20% FBS (Thermo Fisher Scientific – Gibco) and 2% antibiotic-antimycotic (Thermo Fisher Scientific – Gibco). 1.5 mL of this 1% agar/1X DMEM solution was transferred into 35 mm tissue culture dishes and allowed to cool to room temperature. Cells were trypsinized, harvested, counted, and resuspended at a concentration of 5 million cells/mL. 1.4% agarose was melted, cooled to 42°C in a water bath, and 750 μL of the agar was mixed with 650 μL of the 2X DMEM and 100 μL of cell suspension (500,000 cells) to be seeded in each dish. The mixture was added on top of the 1% agar layer and allowed to cool for 30 min before cell culture media was added to the dishes. The dishes were kept in a humidified tissue culture incubator at 37°C and 5% CO2 for three weeks and cell culture media was replenished every three days for the course of the experiment. After three weeks, macroscopic colonies were imaged with the Nikon Eclipse Ti inverted microscope described above using a 20X/0.3 NA A Plan corrected phase contrast objective lens. These large colonies were imaged only at a single focal plane (in which the outermost edges of the colony were most in focus) and images were captured using the Large Image tiling function in NIS Elements software. Individual macroscopic colonies were picked from soft agar using a sterile 200 μL pipette tip and placed in sterile 1.5 mL Eppendorf tubes with 1 mL trypsin. Tubes were placed at 37°C in an incubator for 20 min with occasional pipetting to disrupt the colony. The tubes were then centrifuged for 5 min at 1000 rpm, supernatant was carefully aspirated, cells were resuspended in 1 mL cell culture media, and transferred to a 35 mm tissue culture dish containing another 1 mL of media. Cells were allowed to adhere overnight before preparation for chromosome spreads, following the protocol described above.

### Statistical analysis

Kullback-Leibler (KL) divergence and L2 norm values were determined for each WGD group of each tumor type in order to obtain a measure of the deviation of the aneuploidy landscapes from a uniform distribution. KL divergence was computed using the ‘KL.plugin’ function of the ‘entropy’ package in R, with the event distribution and a uniform distribution as inputs. Similarly, the L2 norm was calculated using the ‘l2norm’ function of the ‘wavethresh’ package in R, with the event distribution and a uniform distribution as inputs.

The significance of the differences in the number of aneuploidies and in the fraction of whole-chromosome aneuploidies between WGD- and WGD+ samples within each tumor type, were determined using a two-sided unpaired Student’s t-test. The significance of the differences in the relative prevalence of the top-3 aneuploidies, the L2 norm values, and the KL-divergence values, between WGD- and WGD+ groups across cancer types, were determined using a two-sided paired Student’s t-test.

Recurrent GISTIC 2.0 events were determined using the p-values and q-values provided by the GISTIC 2.0.23 ‘broad’ analysis output, with the following thresholds used for determining recurrence: q-value<0.05 and recurrence >=0.1. Co-occurrence and mutual-exclusivity of chromosome-arm aneuploidy pairs were determined using the empirical p-values from the permutation analysis (described above), with the following thresholds: p<0.05 and q<0.25.

The number of aneuploidy events of the HCT116 and DLD1 cell lines and their various derivatives, were determined as the sum of whole-chromosome and chromosome-arm aneuploidies in each sample, excluding chromosome Y from the analysis. Karyotypic heterogeneity was determined by counting the fraction of cells with at least one non-clonal whole-chromosome aneuploidy. The significance of the difference between the number of aneuploidy events was determined using a one-sided one-way ANOVA. The significance of the differences in the fraction of whole-chromosome aneuploidies, in the fraction of cells with non-clonal aneuploidies, and in the fraction of gains/losses, were determined using a one-sided Fisher’s Exact test. The significance of the difference in chromosome count distributions between day 0 and day 12 of each WGD group, was calculated using a two-sided Mann-Whitney test.

### Data visualization

Boxplots were plotted using R Base graphics, with the following parameters: bar, median; box, 25^th^ and 75^th^ percentile; whiskers, 1.5 X interquartile range. Heatmaps and histograms were plotted using the ‘ggplot2’ R package, or the ‘matplotlib’ and ‘seaborn’ packages in Python. Venn diagrams were plotted using ‘matplotlib_venn2’ in Python. Whenever data are presented separately for each tumor type, plots were generated only for the 11 tumor types with the highest number of samples, with each WGD group consisting of at least 100 tumor samples.

## Supporting information

Fig. S2 part I

Fig. S2 part II

## Data and Code Access

All datasets are available within the article and its Supplementary Tables, or from the Corresponding Author upon request. The code used to analyze the data is available at https://github.com/kav019/WGD_Aneuploidy.

## Competing interest statement

G.R. is an employee of F. Hoffman-La Roche AG. The other authors declare no competing interests.

## Acknowledgements

The authors would like to acknowledge Jonathan M. Herrera, Eli Reuveni and Gali Yanovich-Arad for technical assistance with computational analyses; Ran Elkon and Ron Shamir for helpful advice and for sharing computing resources; A. Chan Yong Jie and Peter Walian for assistance with data transfer; Caroline Joseph for technical assistance with image file formatting; Sharon Tsach for assistance with figure preparation; and Peter Duesberg for allowing us to collect mFISH data in his lab. This work was funded by the Israel Cancer Research Fund Gesher Award (U.B.-D.), the Azrieli Faculty Fellowship (U.B.-D), the Israel Science Foundation (grant # 1339/18; H.L.), the Virginia Tech College of Science Dean’s Discovery Fund (D.C.), the Virginia Tech ICTAS Center for Engineered Health seed funds (D.C.), the U.S. National Institutes of Health (grant # R01GM140042; D.C.), and Deutsche Forschungsgemeinschaft (STO918-7; Z.S.). Work in the G.R. lab is supported by the Singapore National Research Foundation grant NRFI05-2019-0008.

## Figure and Table Legends

**Supplementary Figure 1: The prevalence and general features of aneuploidy in WGD- and WGD+ tumors (related to Figure 1)**.

(**a**) Comparison of the number of aneuploidies that are significantly (p<0.05, q<0.25) more prevalent in one WGD group than in the other group. (**b**) Comparison of the deviation of chromosome-arm gains (left) and losses (right) from a uniform distribution, between WGD- and WGD+ tumors, across the 22 tumor types. ****, p=5e-08 and p=2e-07, for gains and losses, respectively; two-tailed paired Student’s paired t-test. (**c**) The relative prevalence of chromosome-arm gains (left two columns) and losses (right two columns) in WGD- and WGD+ tumors of 10 selected tumor types. Tumor type abbreviations as in **Fig. 1**.

**Supplementary Figure 2: WGD is associated with significant changes in aneuploidy recurrence patterns (related to Figure 2).**

(**a**) Venn diagrams of the number of the significant (GISTIC 2.0 q<0.05) chromosome-arm aneuploidies in WGD- and WGD+ tumors of 10 selected tumor types. (**b**) Heatmaps of the significance (-log(empirical p-value)) of positive genetic interactions (co-occurrence; in purple) and negative genetic interactions (mutual-exclusivity; in green) between chromosome arms of different chromosomes in WGD- tumors (top two rows) and WGD+ tumors (bottom two rows) of 10 selected tumor types. Events that were significantly co-occurring in one WGD group but significantly mutually exclusive in the other WGD group of the same tumor type (empirical p<0.05, q<0.25) are highlighted on the heatmaps. (**c**) Venn diagrams of the number of the significantly co-occurring (empirical p<0.05, q<0.25) chromosome-arm inter-chromosomal genetic interactions in WGD- and WGD+ tumors of 10 selected tumor types. (**d**) Histograms of the distribution of enrichment scores (defined as -log(empirical p-value)) for co-occurrence (left) and mutual exclusivity (right) of inter-chromosomal genetic interactions in WGD- vs. WGD+ tumors of BRCA, GBM, LGG and LUAD tumors. Discordant genetic interactions, which are cooccurring in one of the WGD groups but mutually exclusive in the other, are highlighted on the histograms. (**e**) Heatmaps of the significance (-log(empirical p-value)) of positive genetic interactions (cooccurrence; in purple) and negative genetic interactions (mutual-exclusivity; in green) between chromosome arms of the same chromosome in WGD- tumors (left) and WGD+ tumors (right) of 10 selected tumor types. Events that were significantly co-occurring in one WGD group but significantly mutually exclusive in the other WGD group of the same tumor type (empirical p<0.05, q<0.25) are highlighted on the heatmaps. Tumor type abbreviations as in **Fig. 1**.

**Supplementary Figure 3: Isogenic WGD- and WGD+ HCT116 cell lines demonstrate a causal effect of WGD on aneuploidy landscapes (related to Figure 4).**

(**a**) scDNAseq-based comparison of the number of aneuploidies between the near-diploid parental HCT116 cells (HCT116-WT), HCT116 cells expressing GFP (HCT116-GFP), and two WGD+ HCT116 clones (HPT1 and HPT2). ***, p<0.001; two-tailed Student’s t-test. (**b**) The number of whole-chromosome gains and losses observed by mFISH, in HCT116 and its derived WGD+ clones (>20 single cells per clone). (**c**) The relative fraction of whole-chromosome aneuploidies relative to arm-level aneuploidies (top) or structural aneuploidies (bottom; including arm-level aneuploidies, translocations and smaller structural alterations), in HCT116 and its derived clones. *, p<0.05, ***, p<0.001; one-tailed Fisher’s Exact test. (**d**) The fraction of cells with whole-chromosome aneuploidies, a measure of karyotypic heterogeneity, in HCT116 and its derived clones. ****, p<0.0001; one-tailed Fisher’s Exact test.

**Supplementary Figure 4: Selection pressures result in non-convergent karyotypic evolution (related to Figure 5).**

(**a**) Bar plots of the number of cells with gains and losses of each chromosome in the near-diploid parental DLD1 cells (DLD1-parental), and 11 DLD1-derived WGD+ clones. (**b**) Representative images of soft-agar macrocolonies emerging from DLD1 cells evolved for 12 days post-cytokinesis failure. Scale, 500 μM. (**c**) Bar plots of the number of cells with gains and losses of each chromosome in the four profiled soft-agar colonies.

**Supplementary Table 1: Summary of the TCGA tumor samples used in this study**. Samples were classified based on their WGD status. The 22 tumor types with > 20 samples in each WGD group were included in the analyses.

**Supplementary Table 2: Results of an arm-level GISTIC 2.0 analysis of the TCGA tumors.** Shown are the full GISTIC 2.0 results for each of the 22 tumor types, including the frequency, frequency scores and q-values for the amplification (gain) and deletion (loss) of each chromosome arm.

**Supplementary Table 3: Inter-chromosomal genetic interactions between aneuploid chromosome-arms**. Shown are the p-values and q-values of all possible inter-chromosomal chromosome-arm genetic interactions. For each of the 10,000 permuted matrices, the significance of co-occurring and mutually-exclusive events was calculated using a hypergeometric test, thereby generating a background distribution of 10,000 (permuted) p-values per each genetic interaction. The observed p-value was compared to the background p-value distribution, and its relative ranking in this distribution was defined as the empirical p-value for that interaction.

**Supplementary Table 4: Intra-chromosomal genetic interactions between aneuploid chromosome-arms**. Shown are the p-values and q-values of all possible intra-chromosomal chromosome-arm genetic interactions. For each of the 10,000 permuted matrices, the significance of co-occurring and mutually-exclusive events was calculated using a hypergeometric test, thereby generating a background distribution of 10,000 (permuted) p-values per each genetic interaction. The observed p-value was compared to the background p-value distribution, and its relative ranking in this distribution was defined as the empirical p-value for that interaction.

**Supplementary Table 5: Summary of the CCLE cell line samples used in this study**. Samples were classified based on their WGD status. The 14 cancer types with >5 samples in each WGD group were included in the analyses.

**Supplementary Table 6: Results of an arm-level GISTIC 2.0 analysis of the CCLE cell lines.** Shown are the full GISTIC 2.0 results for each of the 14 cancer types, including the frequency, frequency scores and q-values for the amplification (gain) and deletion (loss) of each chromosome arm.

