## Supplementary figures and images for "Whole-genome duplication shapes the aneuploidy landscape of human cancers"

### Fig. S1

Supplementary Figure 1

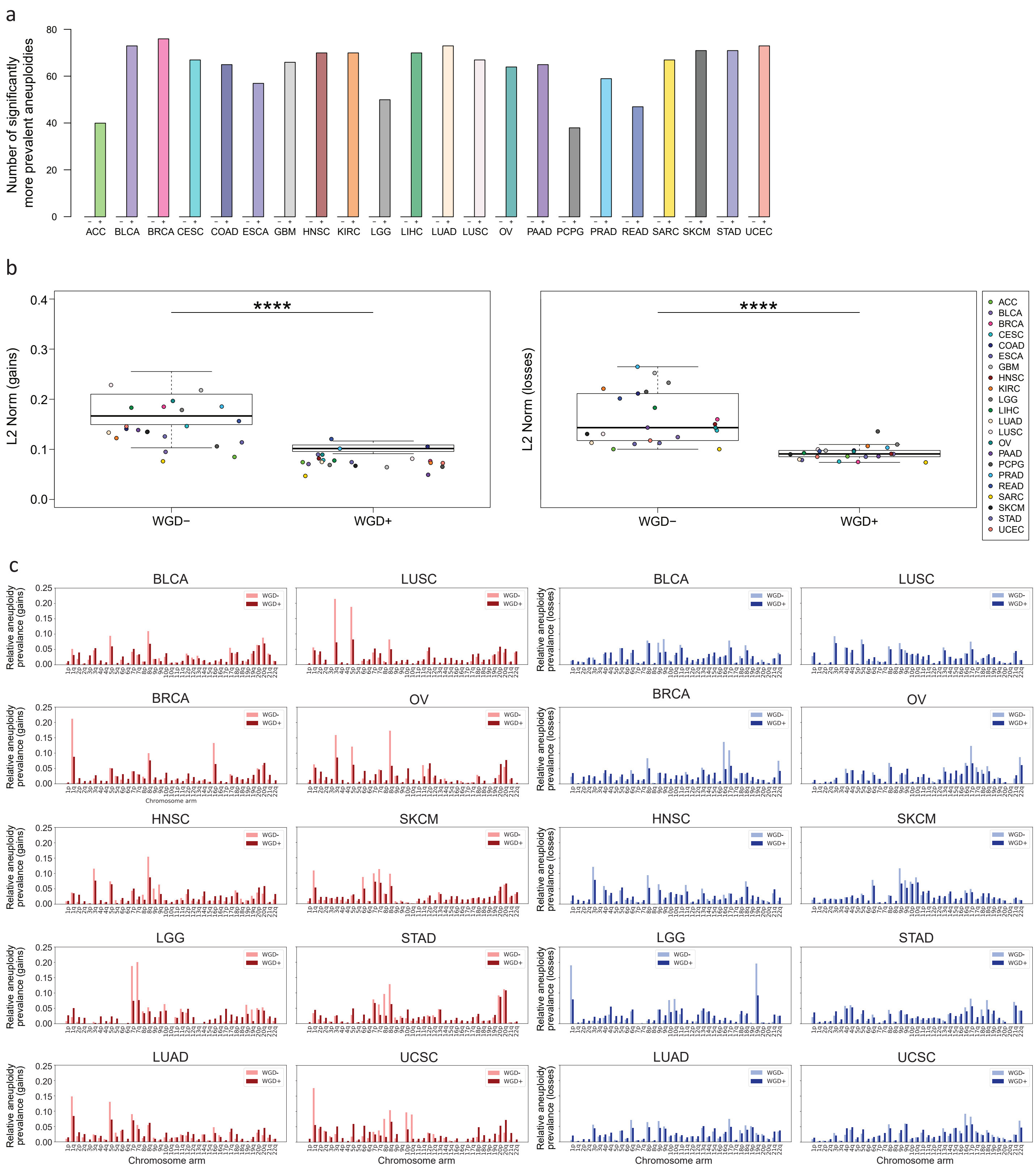

### Fig. S2 part I

Supplementary Figure 2

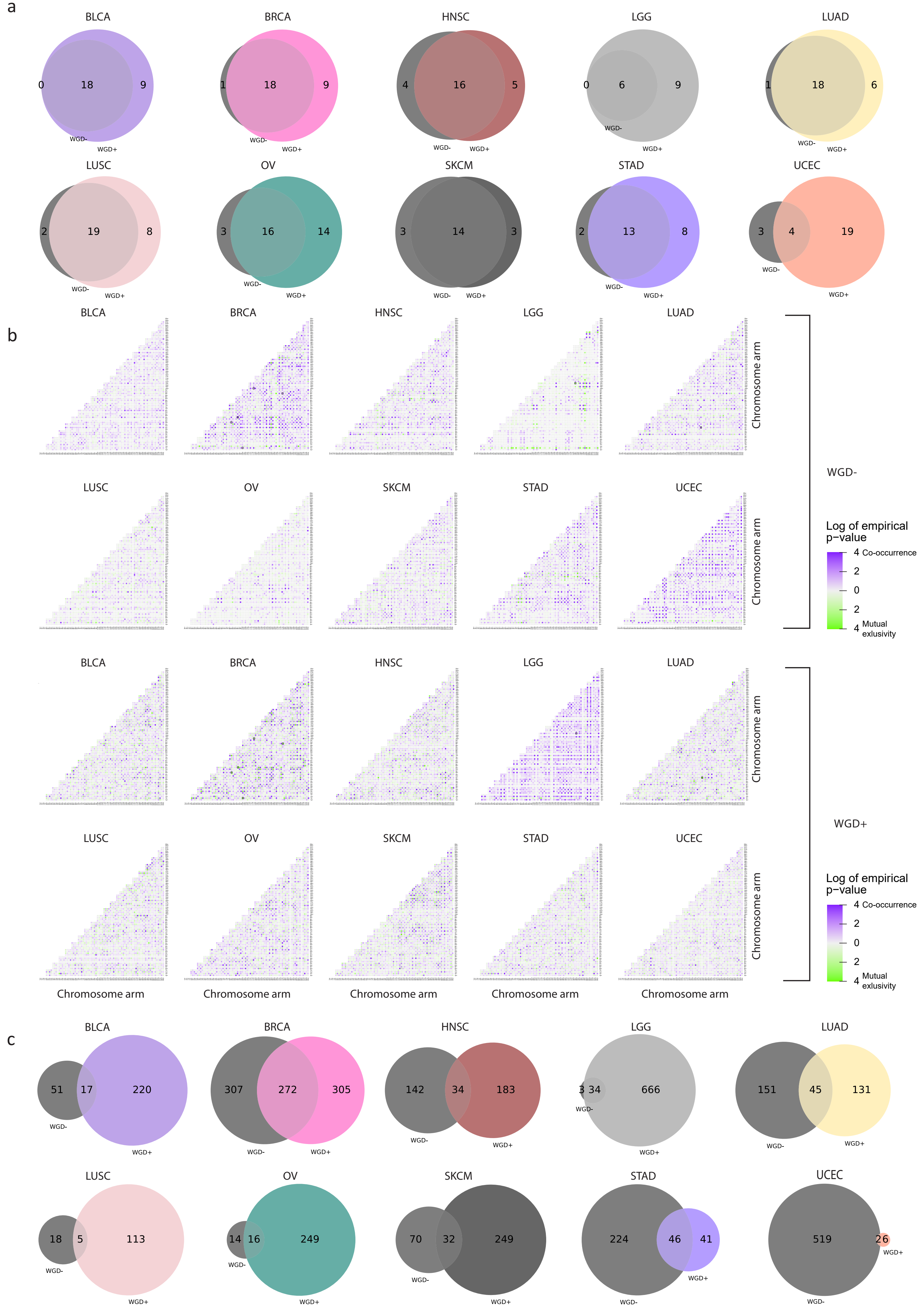

### Fig. S2 part II

#### Supplementary Figure 2 page 2

d

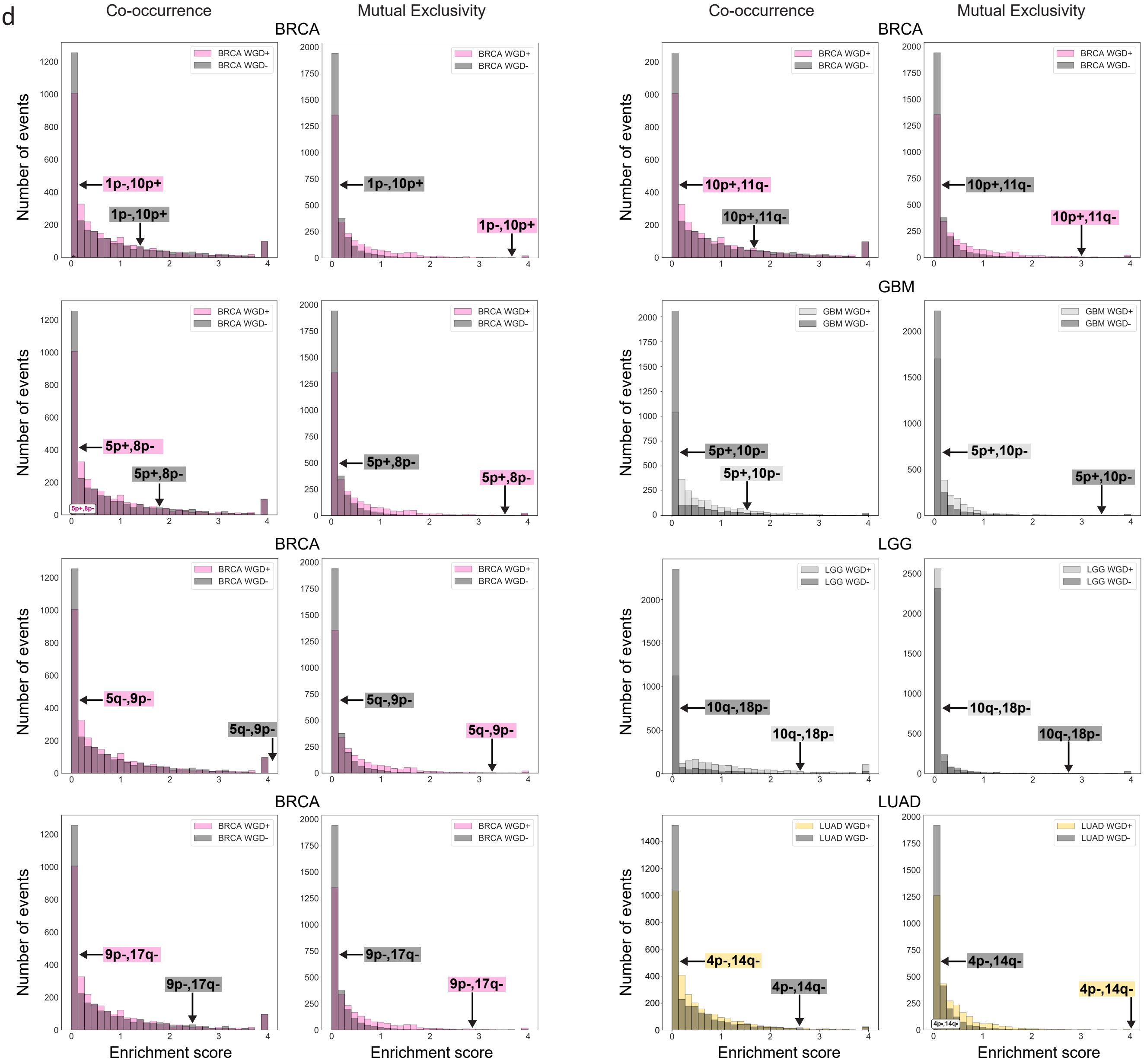


e

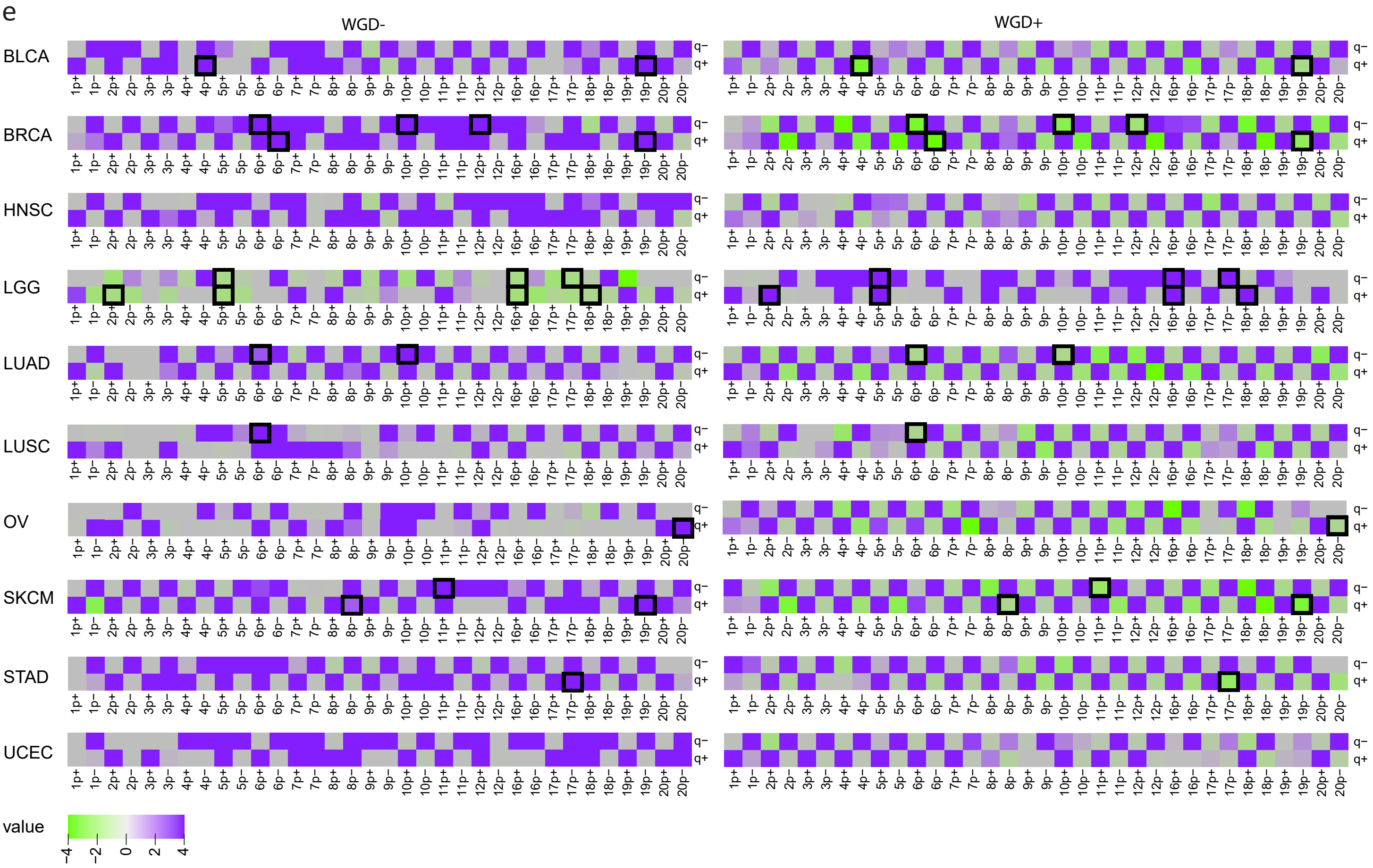

### Fig. S3

Supplementary Figure 3

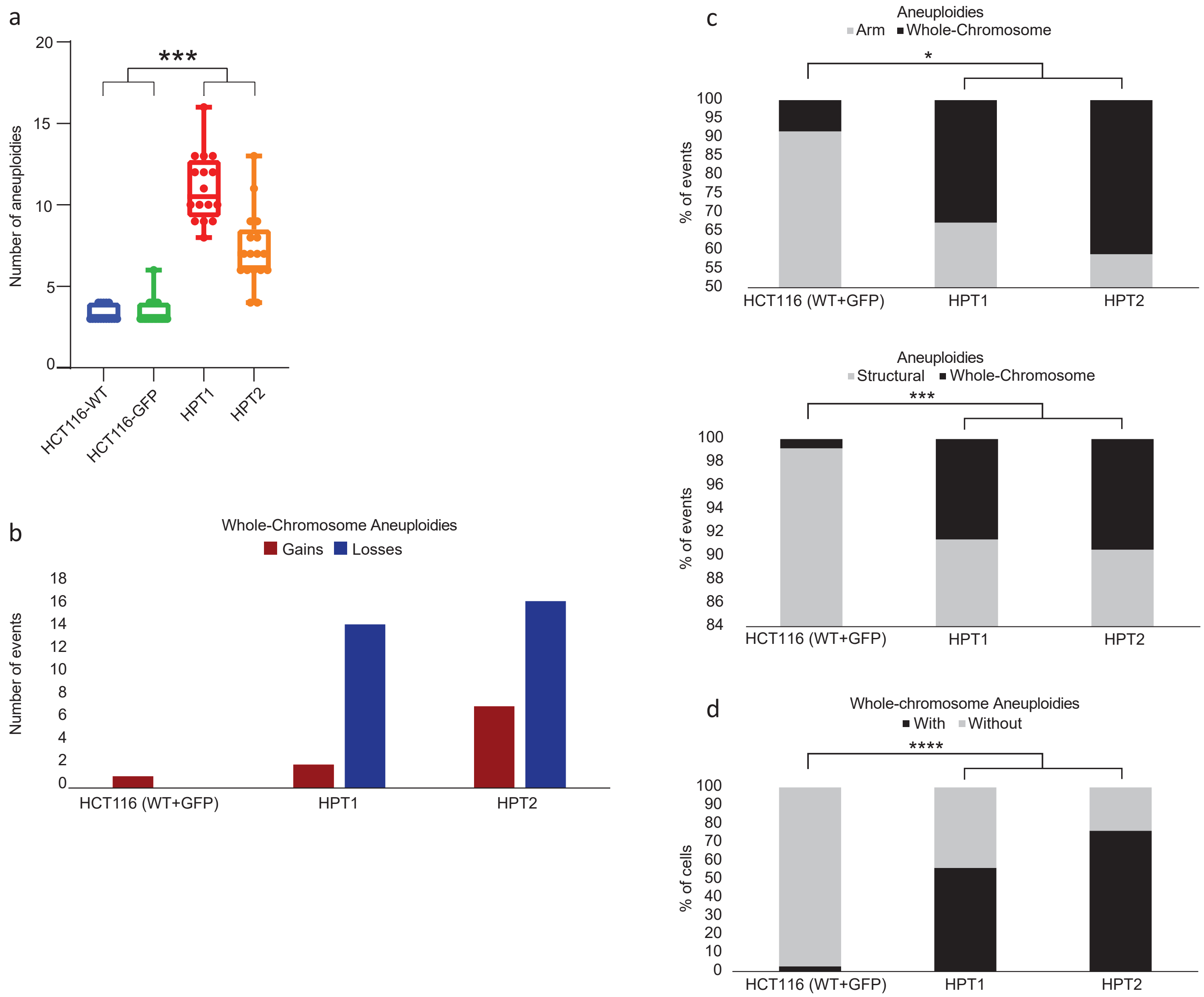

### Fig. S4

#### Supplementary Figure 4

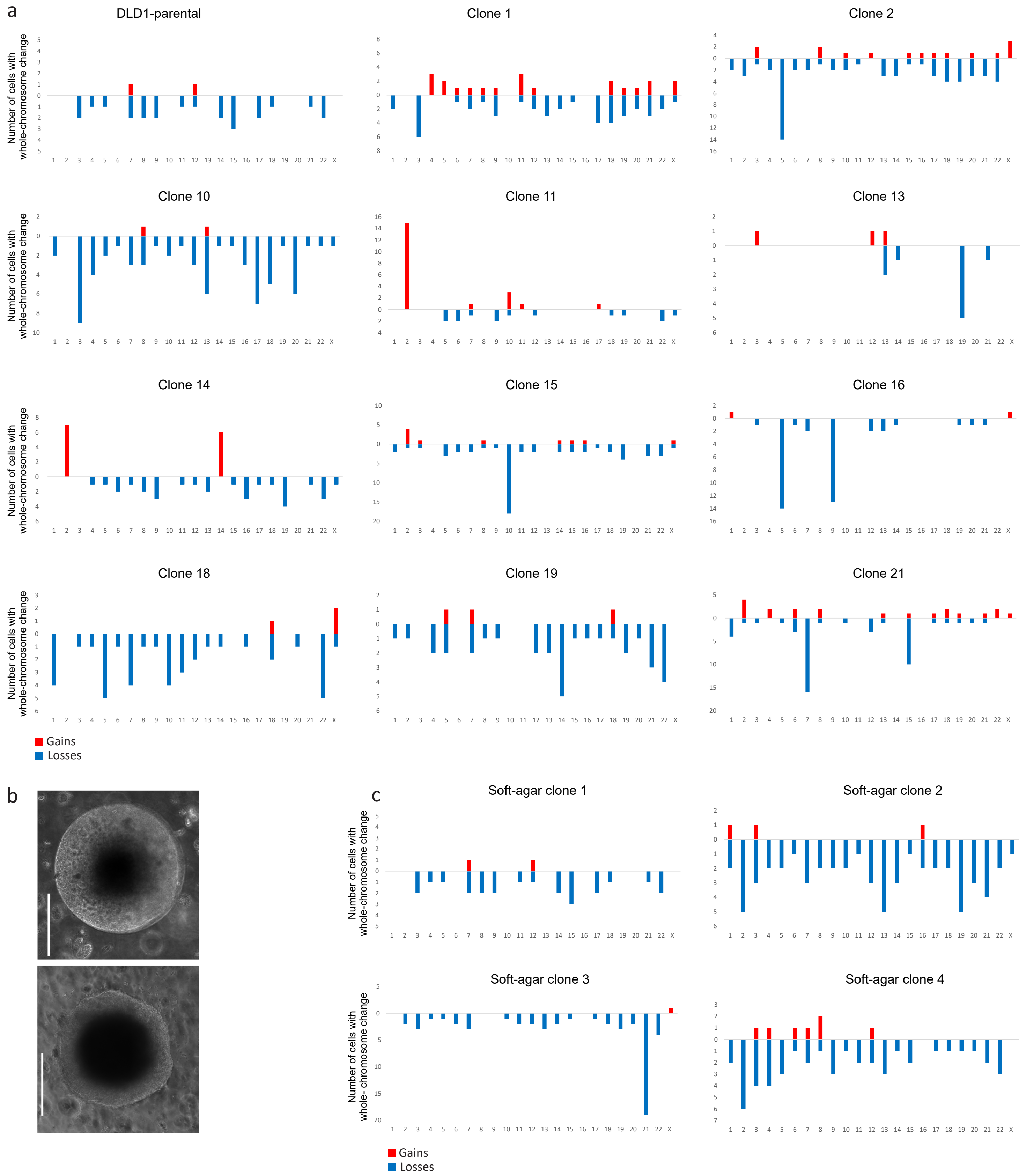
